# Detergent headgroups control TolC folding *in vitro*

**DOI:** 10.1101/2022.04.28.489915

**Authors:** Ayotunde Paul Ikujuni, S. Jimmy Budiardjo, Joanna S.G. Slusky

**Affiliations:** Department of Molecular Biosciences, The University of Kansas, 1200 Sunnyside Ave, Lawrence, KS 66045; Center for Computational Biology, The University of Kansas, 2030 Becker Dr., Lawrence, KS 66045-7534

**Author notes:** These authors contributed equally (Ayotunde Paul Ikujuni, S. Jimmy Budiardjo).

## Abstract

TolC is the trimeric outer membrane component of the efflux pump system in *E. coli* responsible for antibiotic efflux from bacterial cells. Over-expression of efflux pumps has been reported to decrease susceptibility to antibiotics in a variety of bacterial pathogens. Reliable production of membrane proteins allows for the biophysical and structural characterization needed to better understand efflux and for the development of therapeutics. Preparation of recombinant protein for biochemical/structural studies often involves the production of proteins as inclusion body aggregates from which bioactive proteins are recovered. Here we find that the *in vitro* folding of TolC into its functional trimeric state from inclusion bodies is dependent on the headgroup composition of detergent micelles used. Nonionic detergent favors the formation of functional trimeric TolC, whereas zwitterionic detergents induce the formation of a non-native trimeric TolC fold. We also find that nonionic detergents with shorter alkyl lengths facilitate TolC folding. It remains to be seen whether the charges in lipid headgroups have similar effects on membrane insertion and folding in biological systems.

## Introduction

The Gram-negative bacterial cell envelope contains a distinctive extra protective layer called the outer membrane. The outer membrane not only acts as a protective layer to the cell but also functions as a selective permeability barrier controlling the transport of molecules in and out of the cell (1,2). The outer membrane of Gram-negative bacteria is an asymmetric bilayer composed of phospholipids in the inner leaflet, lipopolysaccharide in the outer leaflet, and outer membrane proteins (OMPs) spanning the bilayer (3).

OMPs are almost exclusively antiparallel beta-barrels. They direct diverse cellular functions including signal transduction, general and substrate-specific transport, and enzymatic catalysis (4). OMPs are synthesized in the cytoplasm and targeted to the Sec translocon; a protein complex that helps to transport OMPs across the inner membrane (5). To traverse the periplasmic space, OMPs are aided by periplasmic chaperones (6-9). The folding of the majority of OMPs requires the BAM machinery which catalyzes insertion into the outer membrane (10-14).

For biochemical and structural studies which often require a high concentration of proteins, isolating OMPs from their native environment is extremely labor-intensive. This is because the small volume of the outer membrane results in low-yield protein extraction. When possible, refolding of recombinant protein is preferred.

Preparation of recombinant outer membrane protein for biophysical or structural studies often involves production of proteins as inclusion bodies, aggregates of unfolded, misfolded, and partially folded protein, from which bioactive proteins are recovered (15). When the signal sequence is removed, single chain outer membrane proteins are known to fold from solubilized inclusion bodies into membrane mimetics like detergent micelles (16,17) or lipid bilayers (18,19). When refolding, the selection of detergent or lipid is often chosen empirically or involves screening mimetics with various chemical properties to achieve proper folding.

Previous studies have shown differential effects of various features of membrane mimetics on protein folding. OMP folding rates increase in lipids that form thinner bilayers (20,21), bilayers of increased curvature (21), and bilayers with heat-shock induced lipid dynamics (22,23).

Different types of detergents have been employed in the solubilization of membrane proteins for *in vitro* structural and biophysical studies. Although nonionic detergents are the most commonly used detergent in studying membrane proteins, zwitterionic detergents are also frequently employed. Of the zwitterionic detergents used, LDAO is the most common for biochemical and structural analysis of OMPs (24). OMPs that have been studied with LDAO range from small 8-stranded OMPs, OmpW and Ail (25,26), 12-stranded autotransporter, EstA (27),14-stranded long-chain fatty acid transporter, FadL (28),16-stranded OMPs sortase, BamA (29) to the multi-barrel porin, OmpF, with three 16-stranded barrels, (30)

Outer membrane efflux pumps present a unique challenge in refolding since they are composed of three separate chains. We recently found that with an additional concentration step, multimeric outer membrane proteins such as TolC can be folded into membrane mimetics like single chain barrels are (31). TolC is an outer membrane protein that serves as a component of the tripartite resistance-nodulation-cell-division (RND) efflux pump in *E. coli*, responsible for the expulsion of toxins, including antibiotics, from bacterial cells (32,33). Tripartite RND efflux pumps are composed of three protein subunits that span the inner and outer membranes – where energy from the proton motive force of the inner membrane pushes small hydrophobic molecules out through the periplasmic adapter protein, and then the outer membrane channel (34). The assembly of these three subunits of this pump facilitates extrusion of antibiotics and other toxins from bacterial cells (35). Expression of the AcrABTolC efflux pump is directly correlated with antibiotic resistance in clinical isolates as well as *in vitro* selected mutants of different bacterial pathogens (36-38).

The outer membrane pore of TolC is composed of three identical chains **(Figure 1A)** and the folding of TolC into its native trimeric state is essential for the protein to engage the periplasmic subunit of the efflux pump. Because of the prominent role that TolC plays in efflux pump mediated antibiotic resistance, understanding the factors that promote or inhibit the assembly of functional TolC is important for developing inhibitors of efflux pump assembly.

**Figure 1:**
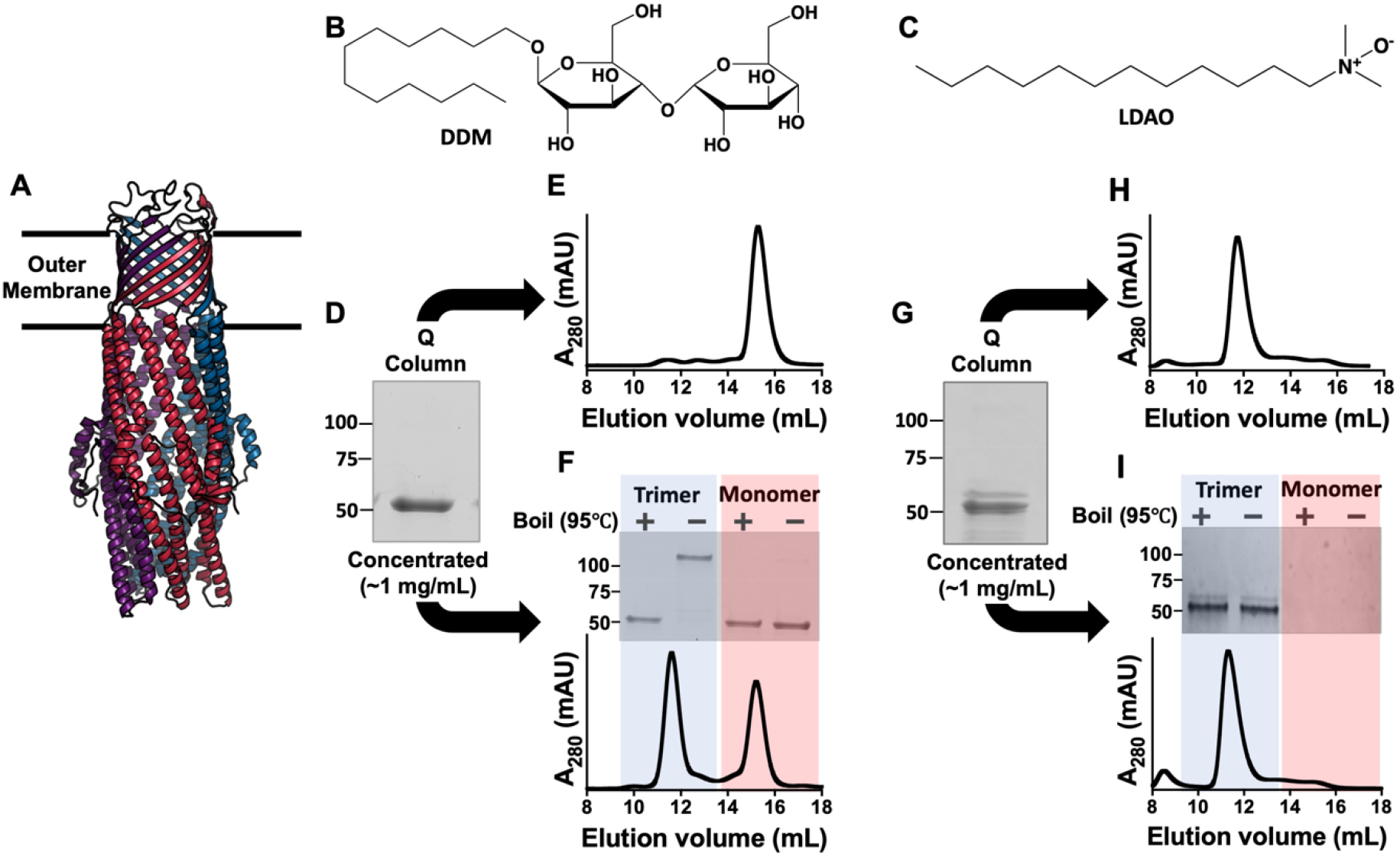
TolC refolded in DDM and LDAO detergents micelles. (A) Structure of trimeric TolC (PDB ID: 1EK9). (B, C) Molecular structure of (A) non-ionic DDM and (B) zwitterionic LDAO. (D) SDS PAGE of TolC in DDM micelles by rapid dilution followed by ion exchange chromatography. (E) SEC chromatogram of TolC elution fraction from ion exchange column in DDM micelles. (F) *Bottom:* SEC chromatogram of concentrated TolC eluate from ion exchange column in DDM micelles. *Top:* SDS PAGE of TolC SEC fractions. (G) SDS PAGE of TolC in LDAO micelles by rapid dilution followed by ion exchange chromatography. (H) SEC chromatogram of TolC elution fraction from Q column in LDAO micelles. (I) *Bottom:* SEC chromatogram of concentrated TolC eluate from ion exchange column in LDAO micelles. *Top:* SDS PAGE of SEC fractions.

In this study, we use *in vitro* refolding experiments and biophysical and biochemical assays to identify a differential effect of detergent headgroup composition on TolC folding *in vitro*. We find that TolC only folds correctly in nonionic detergents and that TolC stability is controlled by the particular nonionic headgroup. In contrast, we find that zwitterionic detergents induce the formation of non-native and non-functional trimeric TolC during *in vitro* refolding from inclusion bodies.

## Results

Different types of detergents have been employed in the refolding and solubilization of membrane proteins for *in vitro* structural and functional studies. Although nonionic detergents like n-dodecyl β-D-maltopyranoside (DDM) are most commonly used detergents for membrane protein purification and structural studies, zwitterionic detergent, n-dodecyl-N,N-dimethylamine-N-oxide (LDAO) is among the top six detergents used for membrane protein purification and crystallization (24). Ionic detergents may be more biologically relevant as phospholipids all have at least one negative charge due to the phosphate in their headgroup. In addition the small micellar size of zwitterionic LDAO is reported to be more favorable for protein crystallization (39). DDM is a nonionic detergent with maltose constituting the headgroup and a 12-carbon alkyl group making up the tail **(Figure 1B)**. In contrast, LDAO is a zwitterionic detergent with an amine oxide headgroup and a 12-carbon alkyl tail (**Figure 1C)**. We have previously reported that *in vitro* refolding of TolC into DDM micelles from inclusion bodies requires three major refolding steps: rapid dilution into detergent micelles, followed by anion exchange chromatography, and then concentration. We found that the third step which involves concentrating the elution fraction from the Q anionic column is essential for the formation of fully folded and functional trimeric TolC (31).

Here we find that the choice of detergent can either cause or prevent TolC folding into its native, functional state. TolC from urea-solubilized inclusion bodies was rapidly diluted into detergent and passed through a Q column. When the detergent was DDM, TolC could then be identified by sodium dodecyl sulfate polyacrylamide gel electrophoresis (SDS PAGE) at approximately 51 kDa, corresponding to the size of a single polypeptide chain of TolC (**Figure 1D)**, consistent with previous reports (31,40,41). When this sample is assessed by size exclusion chromatography (SEC), only a peak corresponding to monomeric TolC (about 15 ml) is visible (**Figure 1E)**, consistent with previous report (31). When concentrated, TolC trimerizes as indicated by the appearance of a second SEC peak at about 11.5 ml **(Figure 1F *Bottom***) and a heat-modifiable, higher molecular weight band on SDS PAGE **(Figure 1F, *Top***).

In contrast, when TolC from urea-solubilized inclusion bodies was diluted into LDAO, the protein is visible at approximately 51 kDa, corresponding to the size of a single polypeptide chain of TolC, on SDS PAGE (**Figure 1G**), and a peak corresponding to the molecular weight of trimeric TolC (about 11.5 ml) is visible by SEC **(Figure 1H**), consistent with previous reports of SEC chromatogram of trimeric TolC (31,42). Upon subsequent concentration, with TolC in LDAO, we find no significant change in SEC (**Figure 1I, *Bottom***) or SDS PAGE (**Figure 1I, *Top***). This indicates that TolC can form a trimeric species, even in the absence of sample concentration. However, this trimeric species may be more unstable as suggested by the dissociation of the trimer on SDS-PAGE even without boiling (**Figure 1I)**.

To better distinguish these two seemingly trimeric states of TolC, we used circular dichroism (CD) spectroscopy to quantify the amount of secondary structure formed in the refolded protein. The mean residue ellipticity (MRE) spectrum reveals the presence of secondary structural elements in TolC refolded in both DDM and LDAO micelles **(Figure 2A)**. The MRE spectrum of TolC refolded in DDM shows two troughs at about 210 nm and 222 nm and a peak at about 192 nm. For TolC refolded in LDAO micelles, the MRE spectrum showed a significantly shallower trough at about 222 nm compared to DDM micelles consistent with less alpha helical content in LDAO. Furthermore, we observed a shift in the trough from ∼210 nm in TolC refolded DDM micelles to ∼206 nm in TolC refolded in LDAO micelles, consistent with the presence of more random coil in LDAO micelles. Also, a general decrease in the signal intensity is observed in the CD spectrum of TolC refolded in LDAO micelles compared to DDM micelles **(Figure 2A)**. The BeStSel (43) webserver was used to estimate the percentage composition of secondary structural elements in TolC refolded in LDAO micelles compared to DDM micelles. The result indicates that TolC refolded in LDAO is less structured **(Figure 2B)**.

**Figure 2:**
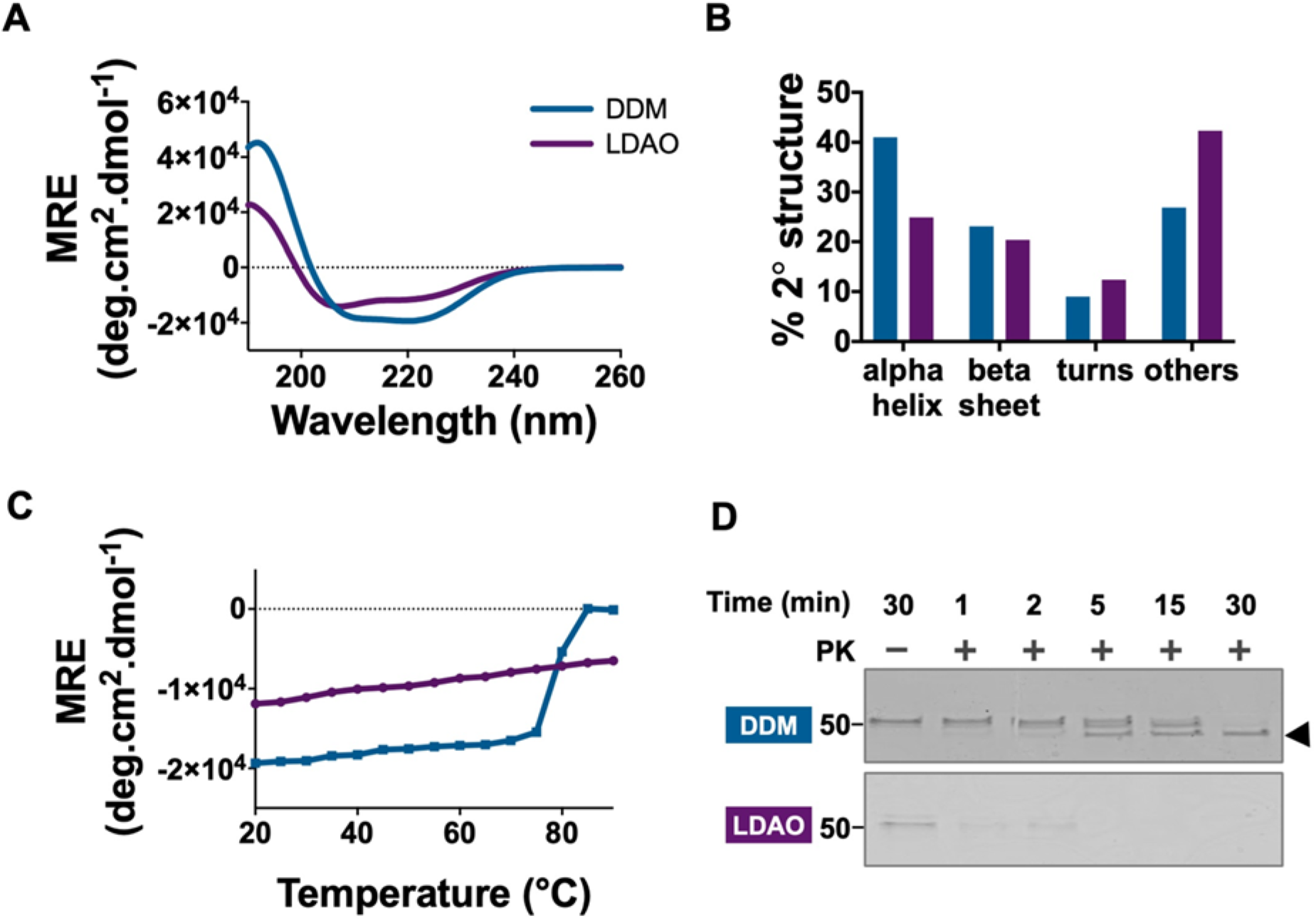
Trimeric TolC is less folded and less inserted in LDAO than in DDM. (A) Overlaid CD spectra of TolC refolded in DDM (blue) and LDAO (purple) micelles. (B) Secondary structure estimation of TolC refolded in DDM (blue) and LDAO (purple) using BeStSel (43) web server. (C) Overlaid mean residue ellipticity (MRE) of TolC refolded in DDM (blue) and LDAO (purple) at 222 nm as a function of temperature. (D) SDS PAGE analysis of trimeric TolC refolded in DDM (blue) and LDAO (purple) incubated with proteinase K. Black arrowhead represents proteinase K digestion resistant product.

To further characterize refolded TolC, we examined the unfolding of the protein by monitoring the change in ellipticity at 222 nm as a function of temperature. For TolC refolded in DDM micelles, we observed a progressive shift in MRE at 222 nm as temperature increased beyond 75 °C, indicating protein unfolding due to high temperature. However, we observed no melting point in the CD signal at 222 nm with TolC refolded in LDAO micelles within the temperature range of 20 to 90 °C used in this study **(Figure 2C and S1)**. These CD data suggest that trimeric TolC formed in LDAO micelles has a different conformation and is possibly less folded than trimeric TolC formed in DDM micelles.

Based on this observation, we used a proteinase K digestion assay to assess the conformation of TolC refolded in LDAO micelles. Previous studies have reported that correctly folded TolC becomes resistant to proteinase K digestion, forming a 46 KDa digestion-resistant product (44,45). Refolded TolC in both DDM and LDAO micelles were subjected to proteinase K digestion. The digestion products were then boiled and visualized by SDS PAGE. TolC refolded in DDM was resistant to proteinase K and formed a 46 kDa digestion resistant product after a 30 minutes incubation. However, TolC refolded in LDAO micelles was completely digested by proteinase K within 30 min of incubation (**Figure 2D**).

The above data indicate that although TolC refolded in LDAO micelles exists as a trimer, it assumes a different conformation than in DDM micelles. We further sought to understand what property of detergents results in the native-like or non-native-like TolC folding. We screened four additional detergents varying headgroups and alkyl chain lengths and observed their effects on the folding, stability, and function of TolC. These include three nonionic detergents n-octyl-β-D-maltopyranoside (OM), n-octylpolyoxyethylene (C8POE), and octaethylene glycol monododecyl ether (C12E8), and one zwitterionic detergent, sulfobetaine 3-12 (SB3-12) **(Table 1)**. Like DDM, C12E8 has a hydrophobic tail with a 12-carbon alkyl group while C8POE and OM have an 8-carbon tail length. The headgroup of DDM and OM is composed of maltose while that of C12E8 and C8POE have polyoxyethylene (**Figure 3A-D**). SB3-12 has a 12-carbon hydrophobic tail length like LDAO, and a positively and negatively charged headgroup contributed by nitrogen and oxygen respectively (**Figure 3D**).

**Table 1:** Detergents used to refold TolC in this study.

|  | Name | Chain length | Headgroup | Headgroup type |
| --- | --- | --- | --- | --- |
| DDM | n-dodecyl-β-D-maltopyranoside | 12 | maltose | Nonionic |
| OM | n-octyl-β-D-maltopyranoside | 8 | maltose | Nonionic |
| C12E8 | octaethylene glycol monododecyl ether | 12 | polyoxyethelene | Nonionic |
| C8POE | n-octylpolyoxyethylene | 8 | polyoxyethelene | Nonionic |
| LDAO | n-Dodecyl dodecyl-N,N-dimethylaminedimethylamine-N-Oxide oxide | 12 | amine oxide | Zwitterionic |
| SB3-12 | sulfobetaine 3-12 (SB3-12) | 12 | sulfobetaine | Zwitterionic |

**Figure 3:**
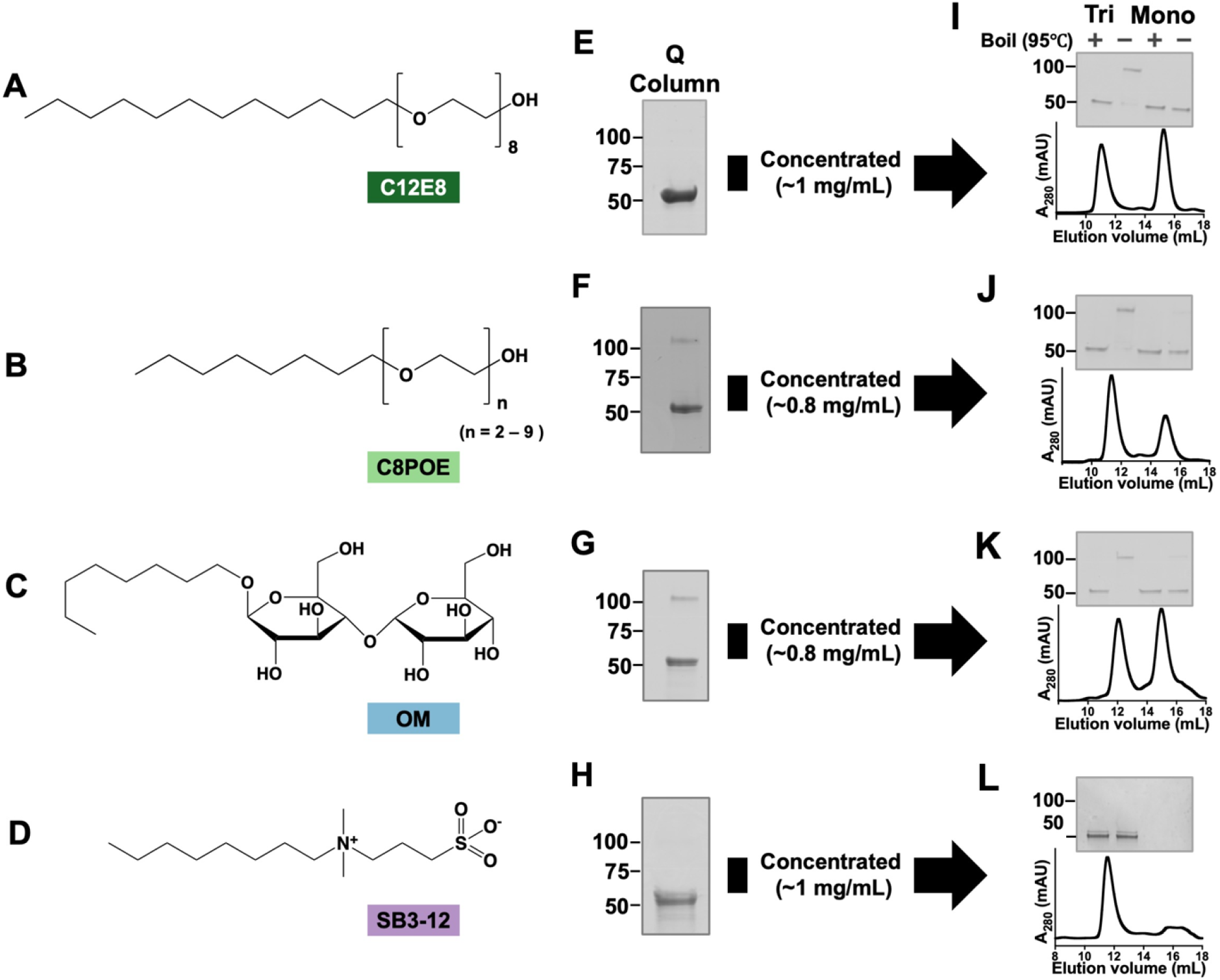
TolC trimerization before concentration depends on detergent headgroup and alkyl chain length. After concentration, TolC trimerization is only dependent on detergent headgroup. **(A-D)** Molecular structures of (A) n-octyl-β-D-maltopyranoside, (B) octaethylene glycol monododecyl ether, (C) sulfobetaine 3-12, (D) n-octylpolyoxyethylene. (E-H) SDS PAGE of TolC refolded in (E) C12E8, (F) C8POE, (G) OM, and (H) SB3-12 micelles by rapid dilution followed by ion exchange chromatography. (I-L) SDS-PAGE and SEC chromatogram of captured monomeric (right) and trimeric (left) TolC refolded in (I) C12E8, (J) C8POE, (K) OM, and (L) SB3-12.

Using an identical protocol to previous TolC folding, urea-solubilized inclusion bodies were rapidly diluted into detergent and then passed through a Q-column. TolC diluted into C12E8 and SB3-12 and visualized on SDS-PAGE, resolved at a molecular weight corresponding to the size of monomeric TolC (**Figure 3E and 3H**). However, SDS PAGE shows that TolC trimerizes more readily in detergent micelles with shorter alkyl chain length C8POE, and OM than longer chain length, DDM and C12E8. (**Figure 3F-G**). However, when concentrated, only the detergents with nonionic head groups developed the trimer band visualized by SDS PAGE which is confirmed with a concomitant trimeric peak on SEC (**Figure 3I, 3J and 3K)**. In contrast, in the zwitterionic SB3-12 detergent micelles, we saw a similar trend observed with zwitterionic LDAO micelles where refolded TolC elutes at a volume corresponding to the size of trimeric TolC on SEC but resolves at a size corresponding to monomeric TolC on SDS PAGE (**Figure 3L**).

We used CD to investigate the folding of TolC in each of the detergents tested. We observed overlaping CD spectra of trimeric TolC refolded in all the nonionic detergents tested (**Figure 4A**) indicating a similar level of secondary structure formation. In the same way, TolC refolded in zwitterionic SB3-12 has a similar CD spectrum to TolC refolded in zwitterionic LDAO (**Figure 4A**). Consistent with TolC refolded into zwitterionic LDAO, TolC refolded into zwitterionic SB3-12, shows no melting point in the temperature range used in this study (**Figure 4B and S1**).

**Figure 4:**
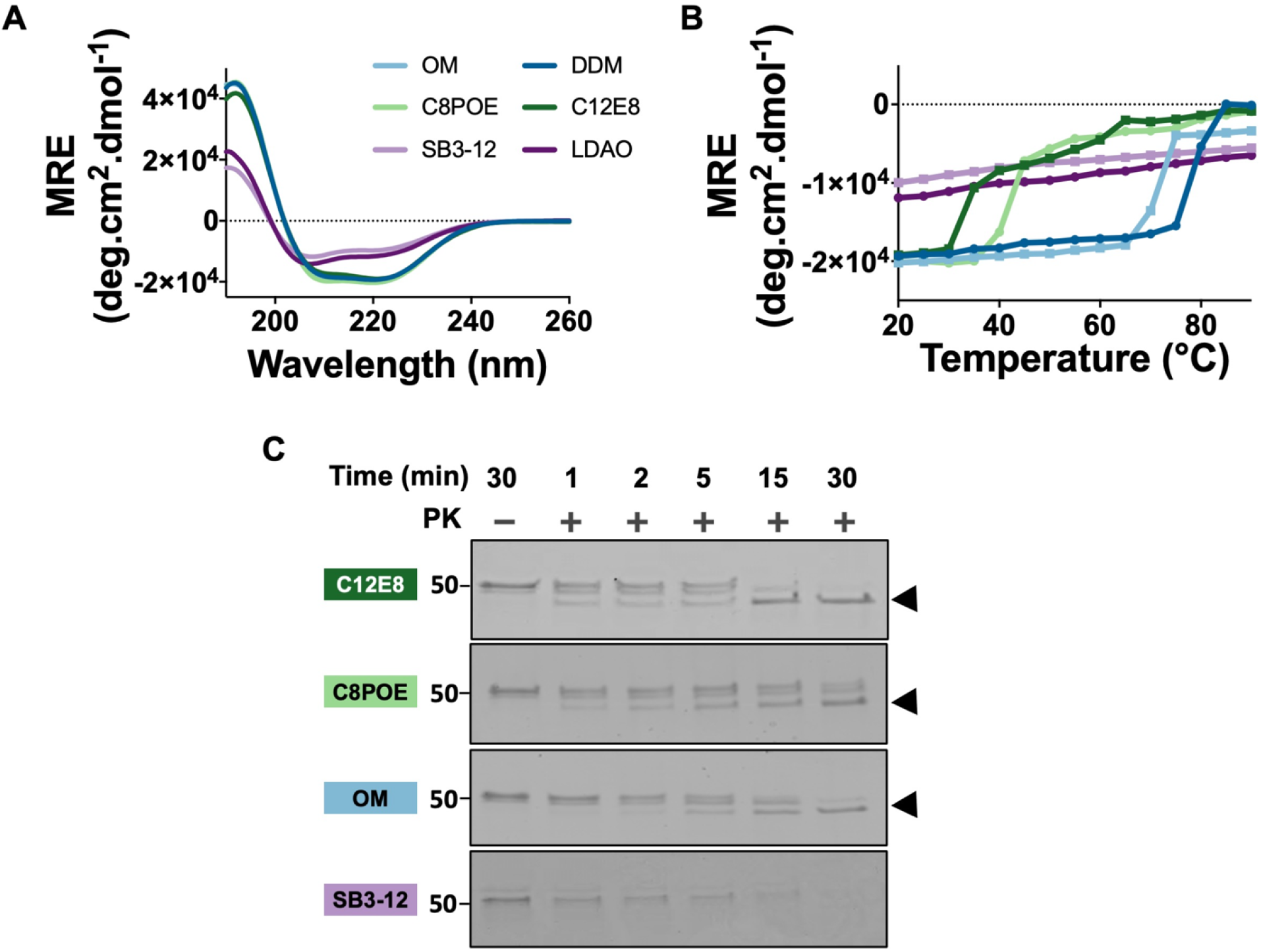
Secondary structure and PK resistance of TolC is dependent on detergent headgroup and not alkyl chain length. (A) Overlaid CD spectra of DDM (dark blue), OM (light blue), C8POE (light green), C12E8 (dark green), LDAO (dark purple), and SB312 (light purple). (B) MRE of TolC at 222 nm as a function of temperature. (C) SDS PAGE analysis of trimeric TolC incubated with proteinase K in detergents.

In addition to the differences in the folding in nonionic and zwitterionic detergents, the thermal denaturation assay reveals another head group-based difference in the thermodynamic stability of TolC within the nonionic detergents. Though TolC folded in nonionic detergents demonstrates heat modifiability on SDS-PAGE and has a similar secondary structure as measured by CD, TolC is substantially more stable in the maltose-headgroup lipids. The T_m_ of TolC in the maltose headgroup detergents is 30 – 40 °C higher than the T_m_ of TolC in polyoxyethylene headgroup detergents (**Figure 4B and S1**).

While trimers form even without concentration for either nonionic eight-carbon chain lipid, the effect of chain length on the stability of TolC in the tested nonionic detergents differs depending on the headgroup composition. For detergents with maltose head group, the 12-carbon length chain increased the T_m_ about 7 °C more than the 8 carbon length. Conversely, while for detergents with polyoxyethylene headgroup, the 8 carbon length chain increased the T_m_ about 3 °C more than the 12 carbon.

The observed variance in the stability of TolC in the different detergents tested, led us to investigate the conformation of TolC and its insertion into the tested micelles by PK digestion assay. The PK digestion assay indicates that, like nonionic DDM, trimeric TolC refolded in all nonionic detergents—C12E8, OM, and C8POE—is resistant to proteinase K digestion and forms a 46 kDa digestion resistant product. However, like zwitterionic LDAO, TolC refolded in zwitterionic SB3-12 detergent micelles was completely digested within 30 min of incubation with PK **(Figure 4C)**. This indicates that in all nonionic detergents we tested, TolC is in a native-like conformation, but that in all zwitterionic detergents we tested, TolC is in a non-native conformation.

Because the melting temperature data and proteinase K digestion data indicate a possible difference in conformation between TolC folded in nonionic detergents and TolC folded in ionic detergents, we sought to determine if TolC refolded in zwitterionic detergents (LDAO and SB3-12) can bind native TolC ligand colicin E1 (46,47). Using a coelution assay, a mixture of refolded TolC and ColE1 was incubated for an hour and then separated on an SEC column. We observed a shift in the elution peak in the SEC chromatogram of the mixture of ColE1 and TolC refolded in all of the four nonionic detergents tested when compared with that of TolC alone. This elution peak shift indicates the presence of a TolC-ColE1 adduct formed from the interaction of TolC with ColE1 **(Figure 5A-D)**. However, the SEC chromatogram shows no shift in the elution volume when ColE1 is mixed with trimeric TolC refolded in LDAO or SB3-12 micelles. (**Figure 5 F-G**). Therefore, although TolC forms a monodispersed trimer when refolded in zwitterionic detergent micelles, the trimer formed has a different conformation from native TolC and does not bind to its endogenous binding partner.

**Figure 5:**
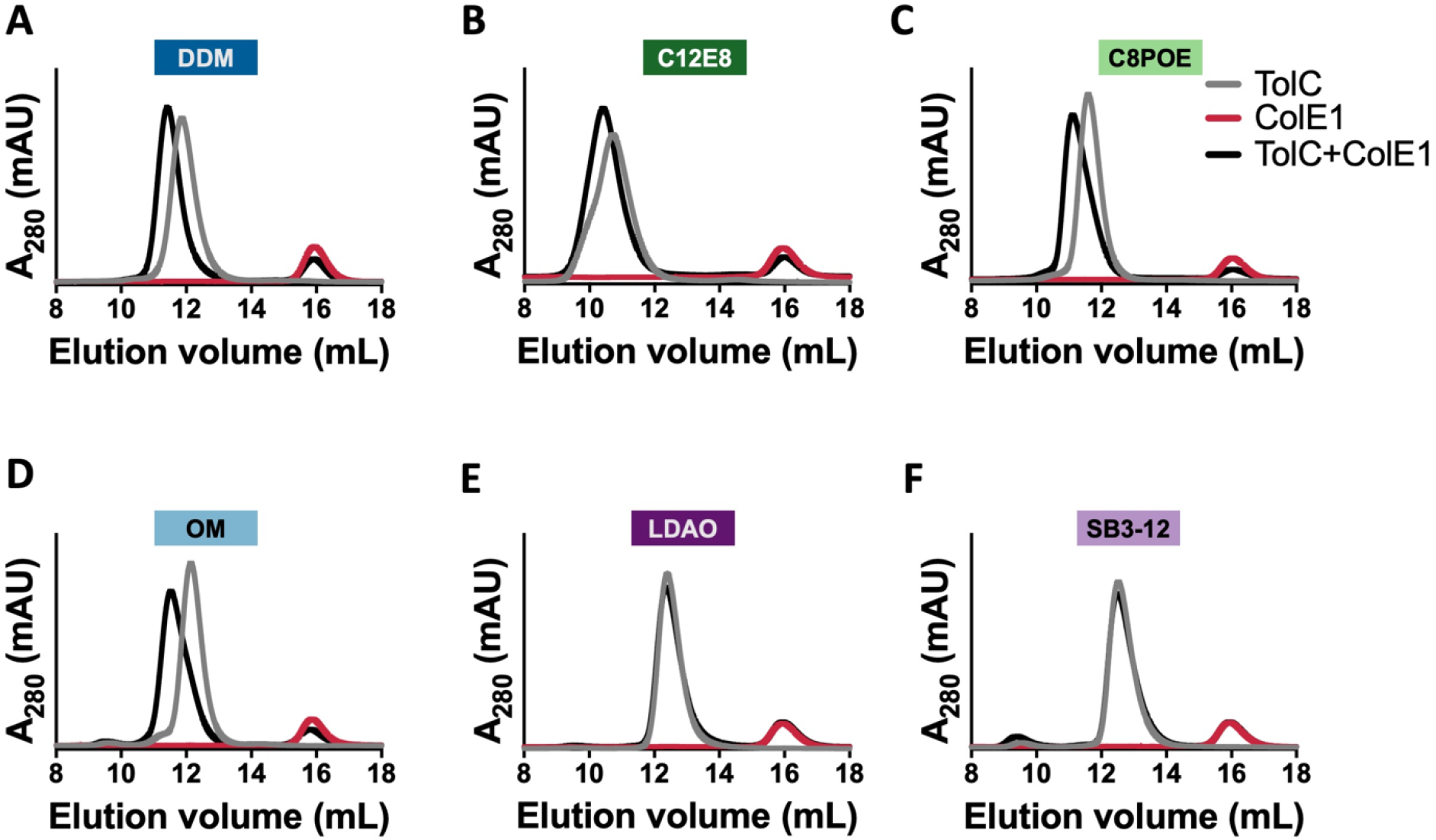
Effect of detergent headgroup on TolC binding to a native ligand. SEC chromatograms of ColE1 (red), TolC+ColE1 (black) with TolC refolded in (A) DDM, (B) C12E8, (C) C8POE, (D) OM, (E) LDAO, and (F) SB3-12. 10 μM of TolC was incubated with 20 μM of ColE1.

## Discussion

TolC exists in its native and functional state in the outer membrane as a trimer with each polypeptide chain contributing to the formation of a cylindrical, β-barrel crowning an α-helical barrel (45,48,49). This conformation is essential for TolC to perform its function in the outer membrane.

It has only recently been determined that TolC can be folded *in vitro* (31). Here we find a different TolC folding behavior in zwitterionic detergent micelles compared to nonionic detergent micelles. Although refolded TolC forms a monodispersed trimer in zwitterionic detergent micelles as observed in the SEC chromatogram **(Figure 1G)**, we find that this is not the native conformation. Structural (**Figure 4)** and functional **(Figure 5)** characterization show that trimeric TolC formed in zwitterionic detergent micelles has a different conformation than native trimeric TolC. Although we may have expected the soluble, periplasmic alpha-helicalportion to form regardless of the detergent used, we find instead that TolC in zwitterionic micelles have half the alpha-helical content but comparable β-sheet content.

Previous studies have indicated that *in vitro* OMP folding involves the formation of an intermediate step where OMPs associate with the membrane mimetics before they are inserted into the micelles or bilayers (50). The hydrophilic, periplasmic alpha helix domain of TolC is composed of clusters of charged residues. We speculate that the non-native folded state we observed in zwitterionic detergent micelles may be a low-energy state formed by interactions between residues of the alpha helix domain of TolC and the charged headgroup of the zwitterionic detergent micelles. If that were the case, charge-charge interactions would lead to the formation of non-native, stable, but non-functional trimeric TolC on the surface of the detergent micelle.

Our finding that more trimer is visible by SDS page when diluting into detergent micelles with shorter acyl chains before concentration **(Figure 3)** is in agreement with previous studies that showed single-chain OMPs fold more readily into lipids with shorter hydrocarbon chains (20,21,23).

Both the polyoxyethelyne detergent micelles and maltoside detergent micelles remain intact upon heating until well above the TolC unfolding transitions (51-53). Since the unfolding is not due to de-micellation, the unfolding temperature is likely a property of the TolC interaction with the headgroup rather than a property of the micelle itself. Maltose headgroup detergents (DDM and OM) stabilize the TolC trimer at higher temperatures. TolC denaturation in maltoside detergents takes place over a narrow temperature range, starting at about 70 °C to 75 °C and complete denaturation is observed at about 75 °C to 80 °C (**Figure 4B, Figure S1**). The spectra of the semi-denatured state appear more like an alpha helix before complete denaturation **(Figure S1)**. However, native-like trimeric TolC is substantially less stable in detergent with polyoxyethylene headgroups (C12E8 and C8POE). For polyoxyethylene headgroup detergents, TolC denaturation starts at about 40 °C and a gradual loss of secondary structure is observed until complete denaturation is reached at about 65 °C. Moreover, as denaturation begins, the spectra appear more like a beta-sheet before complete denaturation is achieved (**Figure S1**). Thus, the alpha helical part of TolC is possibly disrupted by polyoxyethylene headgroups.

Consistent with previous reports for other OMPs (20,21), we find that shorter alkyl chains facilitate TolC insertion even without concentration. However, we find different effects of chain length on stability depending on the headgroup. Polyoxyethylene increases stability for shorter alkyl chains and maltose headgroups decreasing stability for shorter alkyl chains. Consistent with our temperature melt data, perhaps the long polar region of the oxyethylene headgroups disrupts the helical region of TolC with the longer eight oxyethylene disrupting more than POE which has a distribution of two to nine oxyethylene. This is consistent with small amounts of beta structure remaining in the unfolded state of TolC folded in polyoxyethylene micelles (**Figure S1**). When there are no oxyethylene present TolC is more stable in the longer (C12) alkyl chain of DDM than the shorter (C8) alkyl chain of OM.

Finally, we anticipate that our data will help to better understand the biogenesis of outer membrane efflux pumps. Overall, we find that the head group composition of detergents plays a significant role in the folding of multimeric outer membrane efflux pumps *in vitro*, with zwitterionic lipids preventing TolC folding. These findings may contribute answers to long-standing questions of OMP folding: 1) Why can we so easily refold OMPs *in vitro* but *in vivo* folding requires the BAM complex and 2) Once proto-OMPs are in the periplasm, how does nature prevent OMP folding back into the inner membrane phospholipids. At least for TolC, the answer may be that the natural negative charge of the phosphate in all phospholipids as well as the zwitterionic headgroup of the dominant membrane lipid phosphatidylethanolamine (54) prevents folding into both the outer leaflet of the inner membrane and the inner leaflet of the outer membrane. In contrast, the common laboratory use of non-ionic detergents facilitates TolC folding *in vitro*. It will be important to continue to investigate the biological relevance of this effect and to determine how bacterial cells facilitate efflux pump folding in the presence of zwitterionic lipids.

## Materials and Methods

### *E. coli* strains

E. coli BL21(DE3) was used for the expression of TolC as inclusion bodies.

### Expression and purification of Colicin E1

Colicin E1 was expressed and purified as previously described (47). Briefly, residues 1-190 of colicin E1 (colE1-T) were cloned into pET303 containing a C-terminal 6x histidine tag. Plasmids were transformed into BL21(DE3) and plated on LB + agar + 100 μg/ml carbenicillin. A single colony was inoculated into 20 ml of LB + 100 μg/ml carbenicillin and grown at 37 °C with shaking at 250 rpm overnight. The next morning 1L TB broth + 10 mM MgCl_2_ + 100 μg/ml carbenicillin was inoculated with 20 ml of the overnight starter culture and grown at 37 °C with shaking at 250 rpm until OD_600_ reached 1.0 and induced with 1 mM IPTG. The temperature was reduced to 15 °C with shaking at 250 rpm and incubated for 24 hours. Cell pellets were harvested by centrifugation at 4,000 g for 30 minutes at 4 °C. Cell pellets were resuspended in lysis buffer (TBS, 5 mM MgCl_2_, 0.2 mg/ml lysozyme, 5 μg/ml DNase, 1mM PMSF, 20 mM imidazole) at 3 ml/g of cell pellet. Cells were lysed by sonication in an iced water bath (3 min, 2 sec on, 8 sec off, 40 % amplitude, QSonica Q500 with 12.7 mm probe). The cytoplasmic fraction was isolated by centrifugation at 50,400 g at 4 °C for 1 hour. The supernatant was filtered through 0.22 μm membrane filter and applied to a 5 ml HisTrap FF column and purified using an ÄKTA FPLC system with 20 column volumes wash step with binding buffer (20 mM Tris, 40 mM NaCl, 25 mM imidazole) and eluted using a linear gradient from 0 – 100% elution buffer (20 mM Tris, 400 mM NaCl, 500 mM imidazole) in 10 column volumes. Colicin-containing fractions were pooled and concentrated to 2 ml and applied onto a HiLoad 16/60 Superdex 200 pg column and eluted with 1.5 column volumes in 20 mM Tris, 40 NaCl.

### Expression and purification of TolC as inclusion bodies

TolC was expressed and purified as previously described (31). TolC gene was received from R. Misra and cloned into pTrc99a. The signal sequence was deleted for inclusion bodies expression using the forward primer 5’-CATGGTCTGTTTCCTGTGTGAAATTG-3’ and reverse primer 5’-GAGAACCTGATGCAAGTTTATCAGC-3’. A sequence confirmed plasmid of TolC was transformed into *E. coli* BL21(DE3) cells and plated on an LB agar plate with 100 μg/ml of carbenicillin. A single colony from the plate was used to inoculate a 20 ml of LB media containing 100 μg/ml carbenicillin which was then incubated at 37 °C, 250 rpm overnight (∼ 16 h). This starter culture was used to inoculate 1L of LB media containing 100 μg/ml carbenicillin. The culture was grown at 37 °C, 250 rpm until it had an optical density (OD) at 600 nm of ∼0.6. Then expression of TolC was induced with 1 mM of Isopropyl ß-D-1-thiogalactopyranoside (IPTG) while the culture remained incubated at 37 °C, 250 rpm for 4 h. Cells were harvested by centrifugation at 4,000 g for 25 min at 4 °C. Cell pellets were resuspended in lysis buffer (TBS, 5 mM MgCl_2_, 0.2 mg/ml lysozyme, 5 μg/ml DNase, 2 mM phenylmethylsulfonyl fluoride (PMSF), 1 % (v/v) Triton X-100) at 3 ml/g of cell pellet and lysed via sonication (3 min, 3 sec on, 7 sec off, 40% amplitude, QSonica Q500 with 12.7 mm probe) in an iced water bath. The lysate was centrifuged at 4,000 g for 25 min at 4 °C and the pellet was resuspended in 40 ml wash buffer (TBS, 1% (v/v) Triton X-100). The lysate was sonicated on an ice bath for an additional three minutes and the inclusion bodies were recovered by centrifugation at 4,000 g for 25 min at 4 °C. The inclusion bodies pellet was washed by resuspension in 40 ml wash buffer and centrifuged at 4,000 g at 4 °C for 25 min. The inclusion bodies pellet was then resuspended in 20 mM Tris-HCl, pH 8, aliquoted, and stored at −20 °C until further use.

### Refolding TolC

TolC was refolded as previously reported (31). Briefly, inclusion bodies were thawed and centrifuged at 4,000 g for 25 min at 4 °C. The inclusion body pellet was dissolved in urea buffer (20 mM Tris-HCl, pH 8, 8 M urea) and incubated at 37 °C for about 15 min. Solubilized inclusion was then centrifuged at max speed on a table-top centrifuge for 5 min to get rid of insoluble inclusion bodies. To initiate refolding, 4 mg/ml of solubilized TolC inclusion bodies was rapidly diluted by 25-fold into refolding buffer (20 mM Tris-HCl, pH 8.0) containing detergent of interest (0.5 % (w/v) n-dodecyl β-D-maltopyranoside (DDM, Anatrace) or 2 % (w/v) n-Octyl-β-D-maltopyranoside (OM, Anatrace) or 1 % (w/v) n-dodecyl-N,N-dimethylamine-N-Oxide (LDAO, Anatrace) or 1 %(w/v) sulfobetaine 3-12 (SB3-12, Sigma-Aldrich) or 1 % (v/v) n-octylpolyoxyethylene (C8POE, Bachem) or 0.5 % (w/v) octaethylene glycol monododecyl Ether (C12E8, Anatrace) and gently mixed on a rotisserie. The solution was passed through a 0.22 μm membrane filter and loaded onto a 5 ml HiTrap Q HP anion exchange column (GE Healthcare, USA) equilibrated with wash buffer (20 mM Tris-HCl pH 8.0) containing detergent of interest (0.05 % (w/v) DDM or 2 % OM or 0.2 % (w/v) LDAO or 2 %(w/v) SB3-12 or 0.6 % (v/v) C8POE or 0.05 % (w/v) C12E8). The column was then washed with 5 column volumes of wash buffer. The protein was then eluted with 3 column volumes of elution buffer (20 mM potassium phosphate buffer pH 8, 500 mM sodium fluoride) containing the same concentration of detergent of interest in wash buffer. 2X Laemmli SDS sample buffer was added to each sample and loaded on a 4 – 20 % polyacrylamide gel for SDS-PAGE analysis. Gels were imaged with an Epson Perfection V600 photo scanner.

### Size exclusion chromatography

Elution fractions from TolC refolding above were pooled and concentrated using an Amicon centrifugal protein concentrator of 10K molecular-weight cutoff at 4,000 g and 4 °C to a concentration of about 1 mg/ml unless otherwise stated. The sample was then filtered with an 0.22 µm filter. 500 µl of sample was loaded and the loop was emptied with 2.5 ml elution buffer onto a Superdex 200 Increase 10/300 pg size exclusion column, pre-equilibrated with the elution buffer. The sample was then eluted with the elution buffer containing the detergent of interest.

### CD spectroscopy

CD spectra were recorded using a J-815 spectrometer (Jasco, Germany) from 260 to 190 nm. Thermal melt spectra were obtained by CD between 20 °C and 90 °C at 0.5 nm intervals in a 0.1 cm pathlength quartz cuvette. Spectra were collected at 5 °C temperature interval and 5 °C/min temperature ramp time with two minutes of equilibration time at each target temperature. Two replicates were collected for each protein sample and baseline (elution buffer) at 100 nm/min scanning speed, a digital integration time of two seconds, and a bandwidth of 1.5 nm. The spectra were smoothed with a Savitsky-Golay filter, and the baseline was subtracted from each protein sample. Data were converted to mean residue ellipticity and secondary structure analysis was done using BeStSel web server (43).

### Proteinase K digestion

The assembly of refolded TolC was assessed by proteinase K digestion. 15 µg of protein was treated with 15 µg of Proteinase K on ice for 1, 2, 5, 15, and 30 min. 2 mM PMSF was added to inactivate the protease and the reaction was boiled at 95 °C for 5 min. The susceptibility of protein to the proteinase K was determined by SDS PAGE.

### Coelution

TolC and Colicin E1 were both buffer exchanged into coelution buffer (20 mM Tris pH 8.0, 40 mM NaCl, containing detergent of interest) using PD-10 desalting columns. Binding was determined by coelution on a Superdex 200 Increase 10/300 size exclusion column. 1:2 molar ratio (5 μM TolC trimer to 10 μM ColE1) of TolC and ColE1 respectively were mixed and incubated at room temperature for 1 hour.

## Acknowledgment

We thank Rajeev Misra for the pTrc vector containing TolC gene. We gratefully acknowledge NIGMS awards DP2GM128201 to JSGS, NIGMS awards P20 GM103418 and 2K12GM063651 to SJB. We also acknowledge feedback from Rik Dhar, Ryan Feehan, Daniel Montezano, Alex Bowman, and Samuel Lim.

## Supporting Information

**Figure S1:**
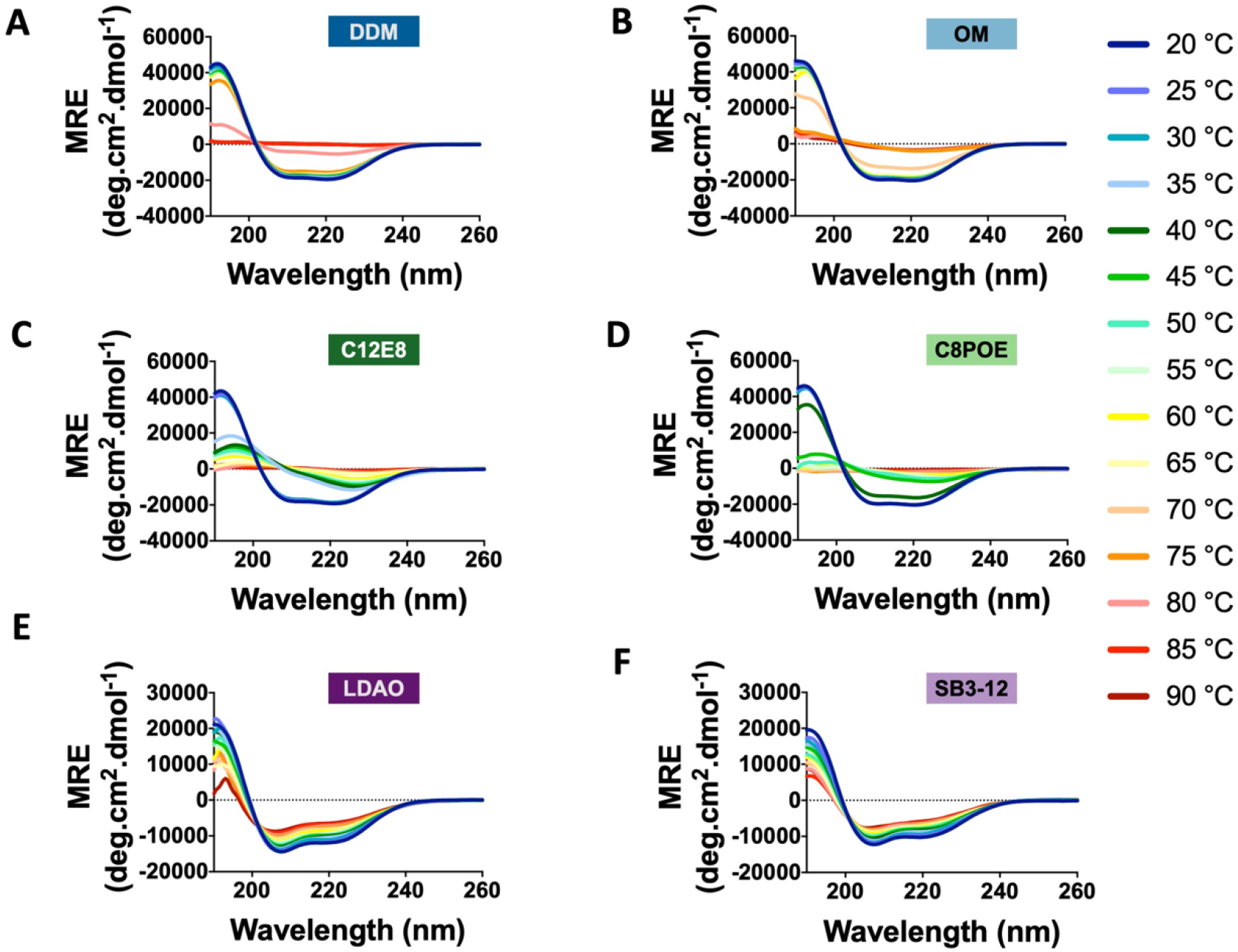
Thermal spectra of TolC refolded in detergents. Change in circular dichroism spectrum per 5 °C increase in temperature from 20 °C to 90 °C of TolC with alpha-helical intermediates in A and B and beta-sheet intermediates in C and D refolded in (A) DDM, (B) OM, (C) C12E8, (D) C8POE, (E) LDAO, and (F) SB3-12.

